# Spiraling Complexity: A Test of the Snowball Effect in a Computational Model of RNA Folding

**DOI:** 10.1101/076232

**Authors:** Ata Kalirad, Ricardo B. R. Azevedo

## Abstract

Genetic incompatibilities can emerge as a by-product of genetic divergence. According to Dobzhansky and Muller, an allele that fixes in one population may be incompatible with an allele at a different locus in another population when the two alleles are brought together in hybrids. Orr showed that the number of Dobzhansky–Muller incompatibilities (DMIs) should accumulate faster than linearly—i.e., snowball—as two lineages diverge. Several studies have attempted to test the snowball effect using data from natural populations. One limitation of these studies is that they have focused on predictions of the Orr model but not on its underlying assumptions. Here we use a computational model of RNA folding to test both predictions and assumptions of the Orr model. Two populations are allowed to evolve in allopatry on a holey fitness landscape. We find that the number of inviable introgressions (an indicator for the number of DMIs) snowballs, but does so more slowly than expected. We show that this pattern is explained, in part, by the fact that DMIs can disappear after they have arisen, contrary to the assumptions of the Orr model. This occurs because DMIs become progressively more complex (i.e., involve alleles at more loci) as a result of later substitutions. We also find that most DMIs involve more than two loci—i.e., they are complex. Reproductive isolation does not snowball because DMIs do not act independently of each other. We conclude that the RNA model supports the central prediction of the Orr model that the number of DMIs snowballs, but challenges other predictions, as well as some of its underlying assumptions.

---

> “[It is not] surprising that the facility of effecting a first cross, the fertility of the hybrids produced, and the capacity of being grafted together… should all run, to a certain extent, parallel with the systematic affinity of the forms which are subjected to experiment…” Darwin (1859)

In the absence of gene flow, the gradual accumulation of divergent genetically based characteristics in different populations can bring new species into being. Some of these divergent characteristics, known as reproductive isolating barriers (Johnson 2006), decrease the level of interbreeding between populations. As populations diverge, isolating barriers accumulate, and the level of reproductive isolation (RI) among populations increases (Coyne and Orr 1989; Roberts and Cohan 1993; Sasa *et al.* 1998; Edmands 2002; Presgraves 2002; Lijtmaer *et al.* 2003; Mendelson 2003; Dettman *et al.* 2003; Moyle *et al.* 2004; Bolnick and Near 2005; Liti *et al.* 2006; Scopece *et al.* 2007; Stelkens *et al.* 2010; Jewell *et al.* 2012; Giraud and Gourbière 2012; Larcombe *et al.* 2015). Eventually RI reaches a point where two of these populations are considered distinct species. Elucidating the precise nature of the relationship between genetic divergence and RI remains one of the central challenges in the study of speciation (Gavrilets 2004; The Marie Curie SPECIATION Network 2012; Nosil and Feder 2012; Seehausen *et al.* 2014).

Dobzhansky (1937) and Muller (1942) proposed a general mechanism through which genetic divergence can cause RI. They noted that, in the absence of gene flow between two populations, an allele that fixes in one population may be incompatible with an allele at a different locus in another population when the two alleles are brought together in hybrids. This negative epistasis, or genetic incompatibility, causes the two populations to become reproductively isolated. Dobzhansky-Muller incompatibilities (DMIs) have been shown to cause inviability or sterility in hybrids between closely related species (reviewed in Presgraves 2010b; Rieseberg and Blackman 2010; Maheshwari and Barbash 2011).

Orr (1995) modeled the accumulation of multiple DMIs as populations diverge. Consider two populations diverged at *k* loci and showing *I_k_* simple DMIs. A *simple* DMI is defined as a negative epistatic interaction between an allele at one locus in one population and an allele at a different locus in the other population. Orr showed that when the next substitution takes place, the expected number of simple DMIs is

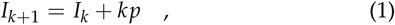

where *p* is the probability that there is a simple DMI between the latest derived allele and one of the *k* alleles at the loci that have previously undergone substitutions (from the population that did not undergo the latest substitution). Assuming *I*_1_ = 0, the solution to difference Equation 1 is

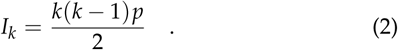

Equation 2 predicts that the number of simple DMIs will accumulate faster than linearly as a function of divergence (prediction #1; Orr 1995). This prediction assumes that *p* remains constant as populations diverge (assumption #1).

DMIs involving *n* ⩾ 3 loci, known as *complex* DMIs (Cabot *et al.* 1994), are also expected to snowball but following different relationships from that in Equation 2: DMIs of order *n* are expected to accumulate at a rate approximately proportional to *k^n^* (prediction #2; Orr 1995; Welch 2004). If DMIs have small, independent effects on RI (assumptions #2 and #3, respectively), then the postzygotic RI they generate is also expected to increase faster than linearly with *k* (prediction #3; Orr 1995). Orr (1995) described this pattern of quantities increasing faster than linearly as “snowballing.” We shall refer to predictions #1–3 of the Orr model collectively as the “snowball effect” (Orr and Turelli 2001).

Several studies have attempted to test the snowball effect. They have employed three different approaches. The first tests prediction #3 of the Orr model: that postzygotic RI snowballs. For example, Larcombe *et al.* (2015) measured the strength of hybrid incompatibility between *Eucalyptus globulus* and 64 species of eucalypts. They observed a faster than linear increase in RI with genetic distance, consistent with prediction #2 of the Orr model. Results from other studies using a similar approach have provided little support for a snowball effect in RI (Sasa *et al.* 1998; Lijtmaer *et al.* 2003; Mendelson *et al.* 2004; Bolnick and Near 2005; Gourbière and Mallet 2010; Stelkens *et al.* 2010; Giraud and Gourbière 2012), leading some to pronounce the snowball “missing” (Johnson 2006; Gourbière and Mallet 2010). However, this approach has several limitations. It can only be applied when postzygotic RI ≪ 1. Furthermore, it only tests one prediction (#3) of the Orr model, and this prediction relies on one assumption (#3) that typically goes untested. Thus, the number of DMIs might snowball (predictions #1–2) even if RI does not.

The second approach tests predictions #1–2 of the Orr model: that the number of DMIs snowballs. For example, Moyle and Nakazato (2010) used a QTL mapping approach to estimate the number of DMIs between species of *Solanum* directly. They introgressed one or a few genomic segments from one species to another. When an introgressed segment caused a reduction in fitness, they concluded that it participated in a DMI. They found that introgressions causing seed sterility accumulated faster than linearly. However, introgressions causing pollen sterility appeared to accumulate linearly, contrary to the snowball effect. Studies following similar approaches have tended to find support for the snowball effect (Matute *et al.* 2010; Moyle and Nakazato 2010; Matute and Gavin-Smyth 2014; Sherman *et al.* 2014; Wang *et al.* 2015).

One advantage of this approach over the first is that it relies on fewer assumptions (#1 compared to #1–3, respectively). However, the second approach also has limitations. The order (*n*) of the DMIs identified is unknown. Therefore, this approach cannot disentangle predictions #1 and #2. Another limitation of these studies is that they are likely to underestimate the true number of DMIs for two reasons. First, the introgressed genomic segments typically contain many genetic differences. For example, the individual segments introgressed in Moyle and Nakazato (2010) included approximately 2–4% of the genome, and likely contained hundreds of genes. Second, individual alleles might participate in multiple DMIs, specially if complex DMIs are common (Guerrero *et al.* 2016).

The third approach tests prediction #1 of the Orr model: that the number of simple DMIs snowballs. Consider two species, 1 and 2, diverged at *k* loci. If an allele, *X*_2_, at one of these loci (*X*) is known to be deleterious in species 1 but is fixed in species 2, then species 2 must carry compensatory alleles at one or more loci (*Y*_2_, *Z*_2_,…) that are not present in species 1 (which carries alleles *Y*_1_, *Z*_1_,… at those loci). In other words, there must be a DMI involving the *X*_2_ and *Y*_1_, *Z*_1_,… alleles.

Following Welch (2004), we define 𝒫_1_ as the proportion of the *k* fixed differences between the species where the allele from one species is deleterious in the other species. For example, Kachroo *et al.* (2015) replaced 414 essential genes of the yeast *Saccharomyces cerevisiae* with their human orthologs. Over half of the human genes (𝒫_1_ = 57%) could not functionally replace their yeast counterparts.

Welch (2004) has argued that estimates of 𝒫_1_ can be used to test the Orr model if two additional conditions are met. If each allele participates in at most one DMI, then we have 𝒫_1_ = *I_k_*/*k*. If, in addition, 𝒫_1_ is entirely based on simple DMIs, then it is expected to increase linearly with genetic distance according to the Orr model (Equation 2)

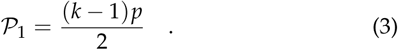

Interestingly, 𝒫_1_ can be estimated without studying hybrids directly. Kondrashov *et al.* (2002) and Kulathinal *et al.* (2004) estimated 𝒫_1_ in mammals and insects, respectively. Surprisingly, both studies reported that 𝒫_1_≈10% and is constant over broad ranges of genetic distances (e.g., human compared to either nonhuman primates or fishes, Kondrashov *et al.* 2002). These results are inconsistent with prediction #1 of the Orr model (Welch 2004; Fraïsse *et al.* 2016). The results of the second and third approaches give inconsistent results, a paradox first noted by Welch (2004). However, the third approach is less direct because it relies on two additional assumptions that have not been tested.

One common limitation to all approaches is that they focus on testing predictions of the Orr model, without testing its assumptions (e.g., assumption #1, constant *p*). Here we use a computational model of RNA folding (Schuster *et al.* 1994; Lorenz *et al.* 2011) to test both predictions and assumptions of the Orr model. The RNA folding model makes satisfactory predictionsof the secondary structures of real RNA molecules (Mathews *et al.* 1999; Doshi *et al.* 2004; Lorenz *et al.* 2011) and has been used to study other evolutionary consequences of epistasis, including robustness (van Nimwegen *et al.* 1999; Ancel and Fontana 2000), evolvability (Wagner 2008; Draghi *et al.* 2010), and the rate of neutral substitution (Draghi *et al.* 2011). We model populations evolving in allopatry on a holey fitness landscape (Gavrilets 2004). In his original model, Orr (1995) made no assumptions on either the evolutionary causes of genetic divergence, or the molecular basis of the DMIs arising from this divergence. Thus, Orr’s predictions should be met in our RNA “world.” Our results provide mixed support for the Orr model.

## Materials and Methods

### Genotype and phenotype

The genotype is an RNA sequence. Unless otherwise stated we used sequences with a length of 100 nucleotides. The phenotype is the minimum free-energy secondary structure of the sequence computed using the ViennaRNA package version 2.1.9 (Lorenz *et al.* 2011) with default parameters.

### Fitness

The fitness of RNA sequence i is determined using the function

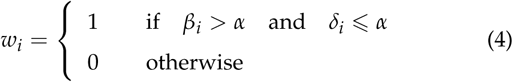

where *β_i_* is the number of base pairs in the secondary structure of sequence *i*, *δ_i_* is the base-pair distance between the structure of sequence *i* and the reference structure, and *α* is an arbitrary threshold. Unless otherwise stated we used *α* = 12. The fitness function in Equation 4 specifies a neutral network (Schuster *et al.* 1994; van Nimwegen *et al.* 1999) or holey fitness landscape (Gavrilets 2004) (Figure 1).

**Figure 1.**
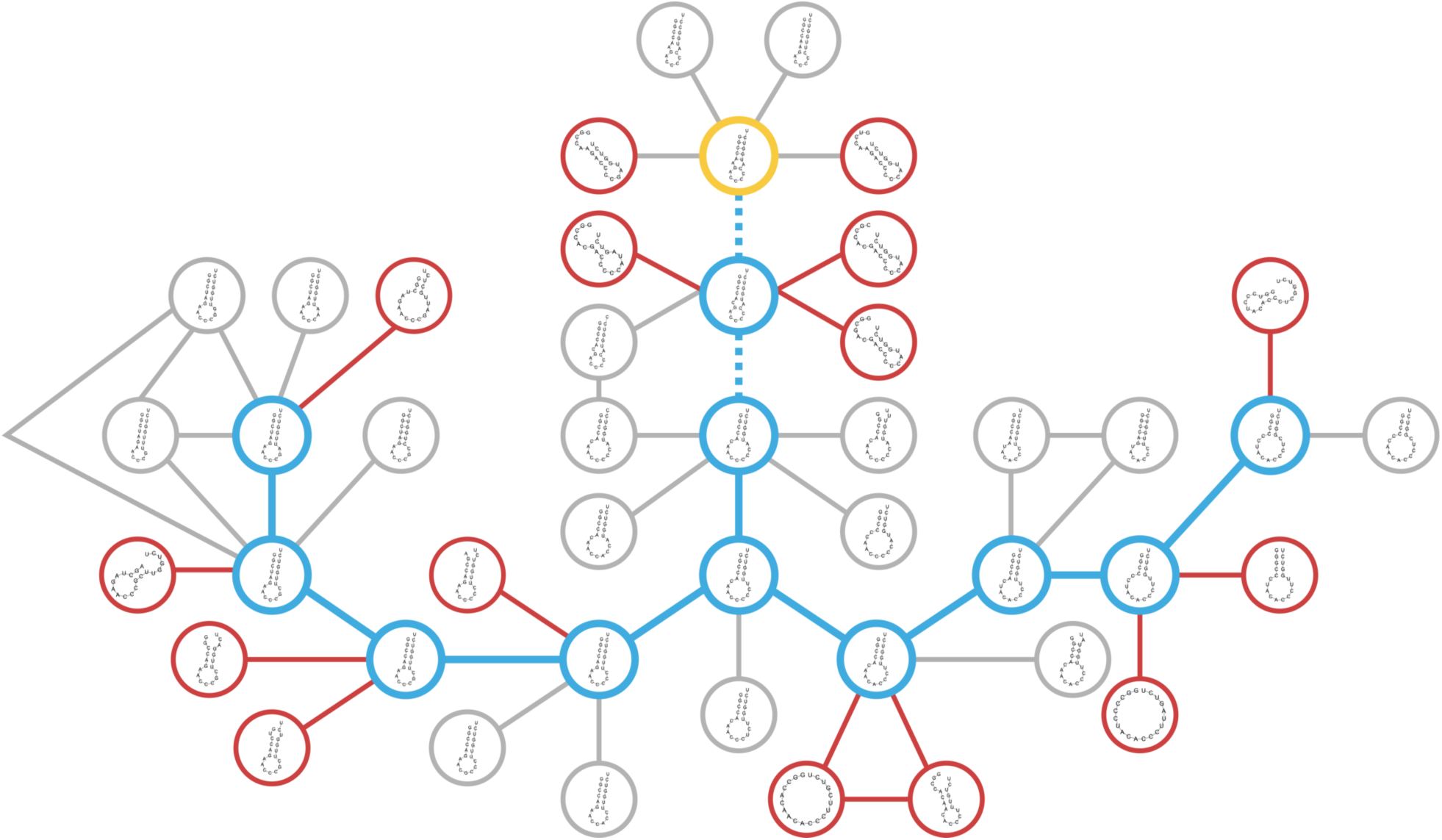
Evolution on a holey fitness landscape. Mutational network of RNA sequences. Lines connect sequences of 20 nucleotides that can be reached by a single nucleotide substitution. Only a tiny fraction of the entire mutational network of ~10^12^ sequences is shown. Furthermore, only a few of the 60 mutational neighbors of each sequence are shown. A sequence is viable (yellow, blue or gray circles) if its secondary structure both has more than *α* = 2 base pairs and is at most *α* = 2 base pairs away from the reference structure (thick yellow circle); a sequence is inviable otherwise (red circles) (Equation 4). Each simulation starts with a burn-in period where a sequence with the reference structure undergoes 3 neutral substitutions (thick dashed blue lines). After that, the resulting sequence is used as the ancestor of two lineages that alternately accumulate neutral substitutions until they have diverged at *k* = 8 sites (thick solid blue lines).

### Evolution

#### Burn-in period

We begin by picking a random viable RNA sequence, define its secondary structure as the reference, and allow it to accumulate 200 random neutral substitutions sequentially, allowing multiple hits. The resulting sequence is used as the ancestor. Table S1 shows summary statistics for the ancestral sequences for *α* = 12.

The burn-in period is necessary because the initial sequence is not representative for the fitness landscape. For example, it has the reference structure (i.e., *δ_i_* = 0 base pairs), whereas most sequences in the fitness landscape are *δ_i_* ≈ *α* base pairs away from the reference structure (Table S1).

#### Divergence

The ancestor is used to found two identical haploid lineages. The lineages evolve by alternately accumulating a series of neutral substitutions without gene flow (allopatry) until they differ at *k* = 40 sites. At a given step, one of the evolving sequences is subjected to a random mutation. If the mutation is neutral, it is allowed to substitute; if it is deleterious, it is discarded and a new random mutation is tried. The process is repeated until a neutral mutation is found. At the next step, the other evolving lineage is subjected to the same process.

At each step, the only sites that are allowed to mutate are those that have not yet undergone a substitution in either lineage since the lineages have started to diverge from their common ancestor. This constraint implies that no more than two alleles are observed at each site during the course of evolution and that substitutions are irreversible, in agreement with the assumptions of the Orr (1995) model. Relaxing this assumption had no effect on the main results (Figure S1; Table S2). All types of base-substitution mutations have equal probability. Insertions and deletions are not considered.

### Inviable introgressions

Two viable sequences, 1 and 2, differ at *k* sites. To detect DMIs of increasing complexity we conduct introgressions of one, two, or three diverged sites from one sequence to another.

#### Single introgressions

We introgress individual nucleotides at each of the *k* divergent sites from sequence 1 to sequence 2 and count the number of inviable introgressions, 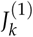. We repeat the procedure in the opposite direction (sequence 2 → 1) and calculate the average of the resulting 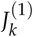 values. The proportion of single introgressions (in one direction) involved in a DMI is given by 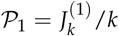 (Welch 2004).

#### Double introgressions

We introgress the *i*(*i* − 1)/2 pairs of nucleotides from sequence 1 to sequence 2, where 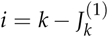 is the number of divergent sites that are not involved in inviable single introgressions in the 1 → 2 direction. We count the number of inviable double introgressions, 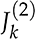 We repeat the procedure in the opposite direction (2 → 1) and calculate the average of the resulting 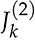 values.

#### Triple introgressions

We introgress all triples of divergent nucleotides from sequence 1 to sequence 2 that contain neither nucleotides involved in inviable single introgressions in the 1 → 2 direction, nor pairs of nucleotides involved in inviable double introgressions in the 1 → 2 direction. We count the number of inviable triple introgressions, 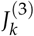 We repeat the procedure in the opposite direction (2 → 1) and calculate the average of the resulting 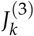 values.

### Potential DMIs

The simple DMIs that might, potentially, affect a sequence can be computed exhaustively by measuring the fitness of all possible single and double mutants derived from the sequence. A potential simple DMI is defined as an inviable double mutant between mutations that are individually neutral.

#### Direct estimation of p

The number of potential simple DMIs, *I*, for a sequence allows us to estimate the value of the parameter in the Orr model (Equation 1) as

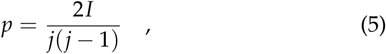

where *j* is the number of neutral mutations tested to estimate *I*.

#### DMI network

We summarize the pattern of interactions between sites using an undirected network where the vertices are sites and the edges represent the existence of at least one potential simple DMI between them (for every pair of sites, there are 9 combinations of double mutants). The resulting network is an example of the networks of interactions described by Orr and Turelli (2001) and Livingstone *et al.* (2012).

#### Reproductive isolation

The degree of RI between two sequences is defined as

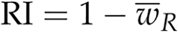

where 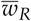 is the mean fitness (Equation 4) of all possible 198 recombinants resulting from a single crossover between the sequences.

#### “Holeyness” of the fitness landscape

For each simulation, we took the ancestor and each of the *k* = 40 genotypes generated during the course of evolution and measured the proportion of their single mutant neighbors (300 per sequence) that are inviable, excluding the 41 original sequences. This proportion estimates the local holeyness of the fitness landscape traversed by the diverging lineages.

### Statistical analyses

All statistical analyses were conducted with R version 3.3.0 (R Core Team 2016).

### Data availability

The software used to run all simulations was written in Python 2.7 and is available athttps://github.com/Kalirad/spiraling_complexity (DOI:http://doi.org/xx.xxxx/zenodo.xxxxxx). The authors state that all data necessary for confirming the conclusions presented in the article are represented fully within the article.

## Results

### Inviable introgressions snowball in the RNA model, but more slowly than expected

The Orr model predicts that DMIs of order *n* should accumulate at a rate approximately proportional to *k^n^*, where *k* is the number of substitutions (prediction #2: Orr 1995; Welch 2004). We tested this prediction using 10^3^ evolutionary simulations with the RNA model. For each simulation, we estimated the numbers of inviable single, double and triple introgressions. We fitted the following model to the average numbers of inviable introgressions at each *k*

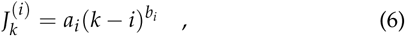

where *i* = 1, 2 or 3 is the number of introgressed alleles, and *a_i_* and *b_i_* are parameters; the *k* − *i* term ensures that 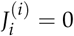

An inviable introgression of *i* alleles indicates the existence of a DMI of order *n* ⩾ *i* + 1. Furthermore, if each inviable introgression is caused by a single DMI, then predictions #1 and #2 of the Orr model lead to the prediction that the exponent *b_i_* ⩾ *i* + 1 (e.g., *b*_1_ ⩾ 2 for inviable single introgressions).

Figure 2 and Table 1 show that inviable introgressions accumulated faster than linearly (*b_i_* > 1) in the RNA model. In addition, the *b_i_* exponent increased with the number of introgressed alleles, *i*, in broad qualitative agreement with predictions #1 and #2 of the Orr model. However, inviable introgressions snowballed much more slowly than expected under the Orr model (*b_i_* ≪ *i* + 2).

**Figure 2.**
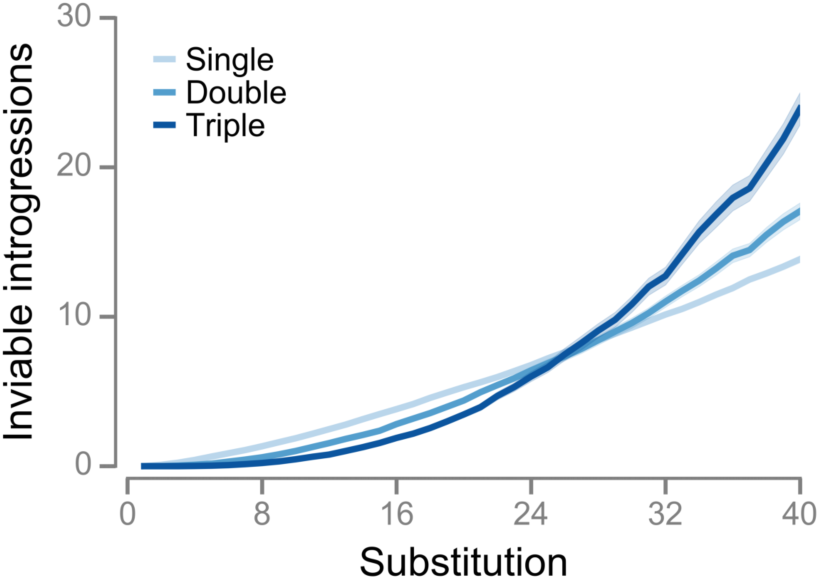
Inviable introgressions snowball in the RNA model, but more slowly than expected. Evolution of the numbers of inviable single, double, and triple introgressions. Values are means of 10^3^ simulations with α = 12. Shaded regions indicate 95% CIs.

**Table 1.** Estimates of the parameters in Equation 6. The model was fitted by nonlinear least-squares regression to the average numbers of inviable single, double, and triple introgressions shown in Figure 2.

| Introgressed alleles, $i$ | DMI order, $n$ | $a_i$ | (95% CI) | $b_i$ | (95% CI) | $R^2$ |
| --- | --- | --- | --- | --- | --- | --- |
| 1 | $\geq 2$ | 0.098 | (0.002) | 1.35 | (0.005) | 0.999 |
| 2 | $\geq 3$ | 0.023 | (0.001) | 1.82 | (0.016) | 0.999 |
| 3 | $\geq 4$ | 0.003 | (0.0004) | 2.44 | (0.032) | 0.999 |

### Some inviable introgressions are caused by multiple DMIs in the RNA model

What explains this mismatch between the RNA data and the Orr model? One possibility is that some inviable introgressions are caused by multiple DMIs (Guerrero *et al.* 2016). For example, imagine two genotypes, 1 and 2, divergent at multiple loci *A*, *B*, *C*,… Genotype *i* has alleles *A_i_*, *B_i_*, *C_i_*,… If there are two simple DMIs between the two genotypes, *A*_1_/*B*_2_ and *A*_1_/*C*_2_, then single introgressions from genotype 1 to genotype 2 would only detect one inviable single introgression.

If we assume that all inviable single introgressions are caused by simple DMIs, then following the Orr (1995) model, the expected number of derived inviable introgressions after *k* + 1 substitutions is given by

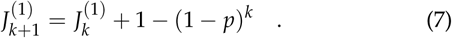

(see File S1 for more details). Assuming 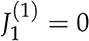, the solution to difference Equation 7 is

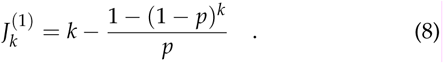

If *p* is low then most inviable single introgressions are caused by single DMIs: 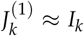. As a result, inviable single introgressions snowball, with *b*_1_ ≈ 2 (Equation 6). As *p* increases, the number of inviable introgressions caused by multiple DMIs increases, and 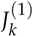 begins to underestimate *I_k_*. In addition, the value of the *b*_1_ parameter in Equation 6 begins to decline, slowing down the snowball as it were. For example, *b*_1_ ≈ 1.92 when *p* = 0.005 (Figure 3A), and b_1_ ≈ 1.61 when p = 0.05 (not shown).

**Figure 3.**
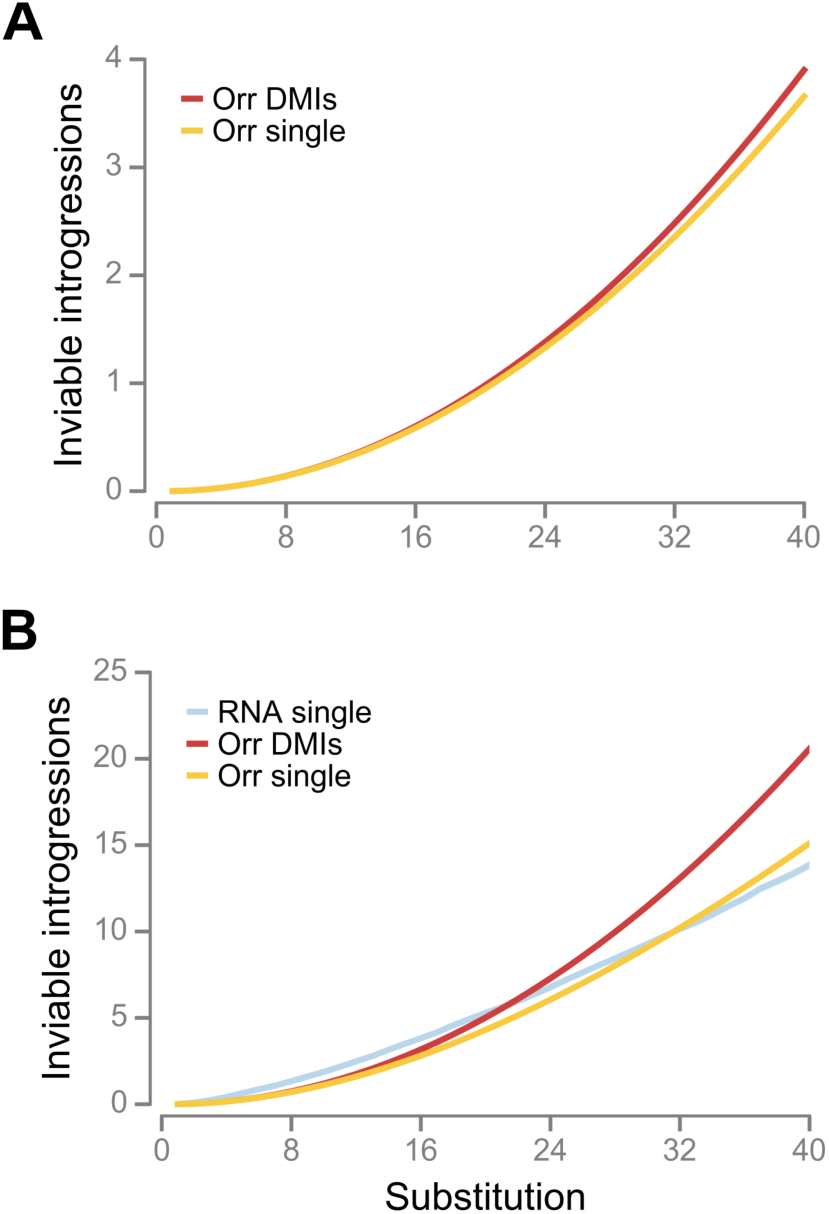
Some inviable introgressions are caused by multiple DMIs in the RNA model. Red lines (“Orr DMIs”) show the evolution of the number of simple DMIs under the Orr model (Equation 2). Yellow lines (“Orr single”) show the evolution of the number of inviable single introgressions assuming that they are all based on simple DMIs evolving according to the Orr model (Equation 8). The blue line in (B) shows the evolution of the number of inviable single introgressions in the RNA model simulations (Figure 2). (A) *p* = 0.005. (B) *p* = 0.0264, obtained by fitting Equation 8 to the RNA data by nonlinear least-squares.

Fitting Equation 8 by nonlinear least-squares regression to the evolutionary response in inviable single introgressions shown in Figure 2 yields an estimate of *p* = 0.0264. If 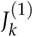 evolved according to Equation 8 with this value of *p*, then *b*_1_ ≈ 1.75 in Equation 6 (Figure 3B), confirming that some inviable single introgressions are caused by multiple DMIs in the RNA model.

However, the lack of agreement between the RNA data and the Orr model is not completely explained by the fact that some inviable introgressions are caused by multiple DMIs for two reasons. First, fitting Equation 6 directly to the same data yields an estimate of *b*_1_ = 1.35, a much lower value than 1.75 (Figure 3B; Table 1). Second, the prediction of *b*_1_≈ 1.75 is conservative because it assumes that all inviable single introgressions are caused by *simple* DMIs. In fact, after *k* = 40 substitutions in the RNA model there were 13.85±0.23 inviable single introgressions, but only 0.600±0.052 simple DMIs (mean and 95% confidence intervals, CIs) (see File S2 for details on how simple DMIs were detected).

### The probability that a simple DMI appears is approximately constant in the RNA model

Another possible explanation the lack of agreement between the RNA data and the Orr model is that p itself has evolved, contrary to assumption #1 of the Orr model. For example, if p declines with divergence according to the relationship *p_k_* = *c*/*k*, where *c* is a positive constant, then simple DMIs are expected to accumulate linearly:

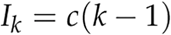

To test assumption #1 of the Orr model, we measured *p* directly in each simulation using Equation 5. We found that *p* declined slightly (Figure S2). However, this decline was not sufficient to explain the estimate of *b*_1_ = 1.35 for the RNA data (Table 1). If 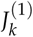 evolves according to Equation 8 with the average value of *p* = 0.0169 estimated from the RNA data, then *b*_1_ ≈ 1.82. If 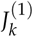 evolves according to Equation 7 with *p* declining with *k* according to the solid line in Figure S2, then *b*_1_≈1.73; *p* would have to show a much steeper decline with *k* to cause *b*_1_ = 1.35 (Figure S2, dashed line).

### Simple DMIs do not persist indefinitely in the RNA model

The direct estimate of *p* in the RNA model shown in Figure S2 reveals another problem for the Orr model. If *p* = 0.0169, then Equation 2 predicts an accumulation of approximately 13 simple DMIs after *k* = 40 substitutions. However, we found that in 57% of runs there were none at all. This discrepancy indicates that a more fundamental assumption of the Orr model may be violated in the RNA model: that simple DMIs, once they have arisen, persist indefinitely (assumption #4). This assumption was not stated explicitly by Orr (1995) and has never, to our knowledge, been called into question.

To test assumption #4, we estimated the DMI networks of sequences as they evolved in our RNA model. Figure 4A shows an example of an RNA sequence evolving on a holey fitness landscape. Initially the sequence displays potential simple DMIs between 21 pairs of sites (Figure 4C). Figure 4B illustrates a potential simple DMI between positions 5 and 12. We refer to these simple DMIs as *potential* because if two diverging lineages each accumulate one of the substitutions underlying one of these DMIs, a simple DMI between the lineages will appear.

**Figure 4.**
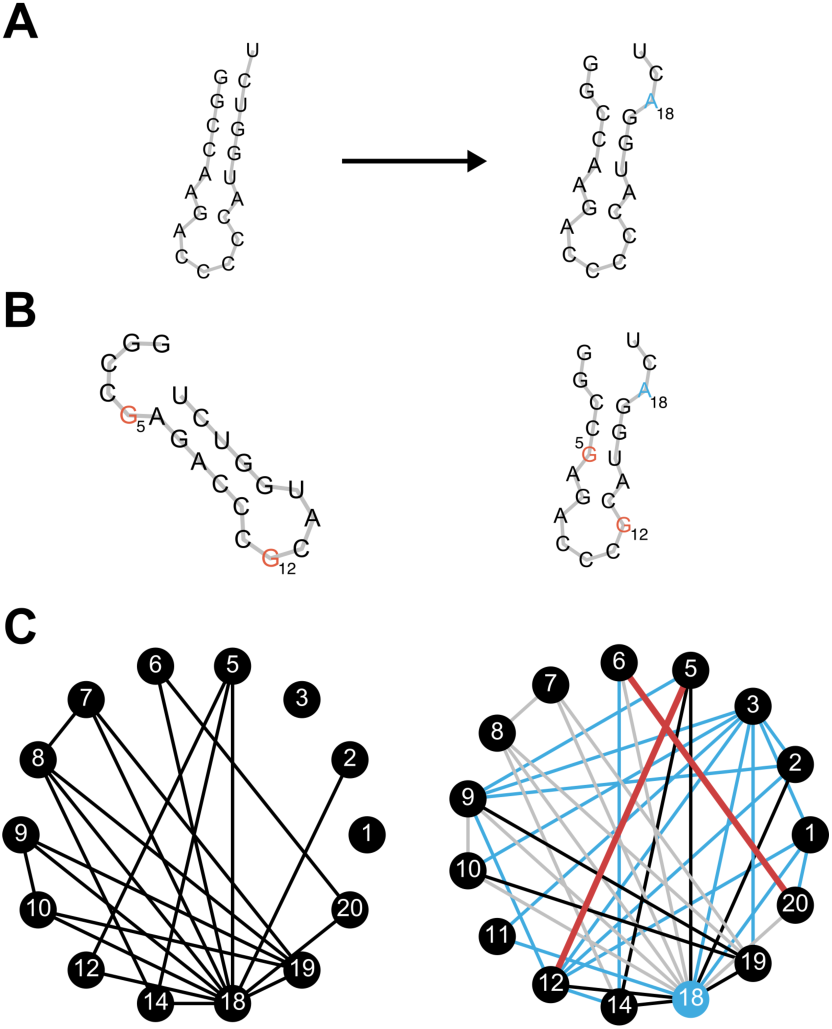
Simple DMIs do not persist indefinitely in the RNA model. (A) The 20 nucleotide long RNA sequence on the left acquires a neutral U→A substitution at position 18 (blue). The holey fitness landscape is defined by *α* = 2 (Equation 4). The secondary structure of the sequence on the left is the reference (*δ*_i_ = 0 base pairs). The structure on the right is *δ_i_* = 2 base pairs away from the reference. (B) There is a potential simple DMI between positions 5 and 12 for the sequence on the left. double mutant at those positions (5: A→G, 12: C→G, red) makes the structure inviable (*δ_i_* = 11 base pairs), even though the single mutations are neutral (not shown). However, a single substitution causes the potential simple DMI to disappear in the sequence on the right, although the single mutations remain neutral in the new background (not shown). In other words, the substitution causes the simple DMI to become complex. (C) A single substitution can dramatically rearrange the network of potential DMIs. DMI networks of the sequences in (A). Vertices correspond to positions in the sequences. An edge in the network on the left indicates that there is at least one potential simple DMI between the two sites (positions 4, 13 and 15–17 have no potential DMIs in either network and are not shown). Black edges in the network on the right are shared between the two networks. Blue edges exist only in the network on the right and indicate the appearance of new potential simple DMIs between sites caused by the substitution. Gray and red edges indicate disappearance of potential simple DMIs in the network on the right. Gray edges indicate disappearances due to the constituent alleles no longer being neutral in the new background. Red edges indicate disappearances caused by complexification; the DMI discussed in (B) is an example (5–12 edge).

The Orr model assumes that the DMI network is static: as populations evolve they actualize potential DMIs (for an alternative, but equivalent, interpretation of DMI networks see Livingstone *et al.* 2012). However, DMI networks are not static in the RNA model. After a single neutral substitution, 13 pairs of sites (62%) lost all potential simple DMIs, and potential DMIs appeared between 18 new pairs of sites (Figure 4C).

The “disappearance” of a potential DMI can occur in one of two ways. First, the substitution may cause the mutations involved in the simple DMIs to become deleterious so that they can no longer participate in potential simple DMIs. A disappearance of this kind means that a potential simple DMI is no longer accessible through independent substitution in two lineages because one of the substitutions cannot take place. Thus, such disappearances do not contradict assumption #4 of the Orr model. The majority of disappearances in Figure 4C (gray lines) are of this kind.

The second kind of disappearance occurs when the substitution modifies the interaction between previously incompatible alleles (red lines in Figure 4C). In other words, the simple DMIs become complex. The potential simple DMI between positions 5 and 12 shown in Figure 4B disappears in this way. This kind of disappearance—complexification—implies that some simple DMIs may not persist indefinitely. In other words, assumption #4 is violated in the RNA model. In the next section we explore the consequences of the complexification of simple DMIs for the snowball effect.

### The modified Orr model

We incorporate the dynamic nature of simple DMIs by extending the Orr (1995) model in Equation 1

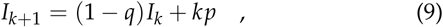

where *q* is the probability that a simple DMI present after *k* substitutions becomes complex after the next substitution. Assuming *I*_1_ = 0, the solution to Equation 9 is

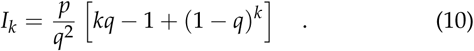

This prediction assumes that both *p* and *q* remain constant as populations diverge.

The original Orr model is a special case of the modified model when *q* = 0. When *q* > 0, the increase in the number of simple DMIs is given by

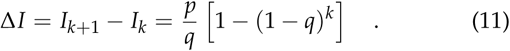

This equation has two consequences. First, the increase in the number of simple DMIs eventually becomes linear with a slope of approximately *p*/*q* when *k* is sufficiently large. Second, if *q* is larger, the “linearization” of Equation 10 occurs for lower values of *k*. Both patterns are illustrated in Figure 5A, which compares the accumulation of simple DMIs under the Orr model with *p* = 0.04 (*q* = 0), and that under the modified Orr model with the same value of *p* and increasing values of *q*.

**Figure 5.**
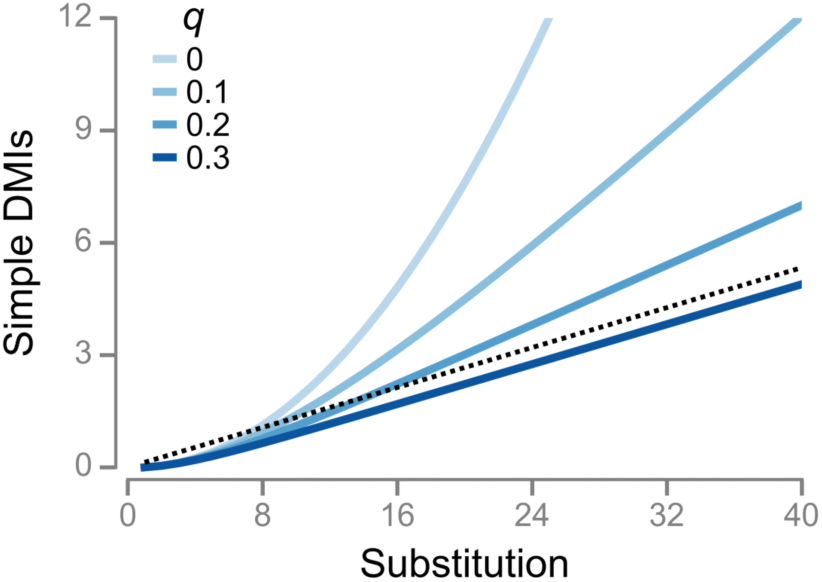
The modified Orr model. Evolution of the number of simple DMIs according to Equation 10. Responses for *p* = 0.04 and different values of *q*. The dashed line shows a slope of *p*/*q* for *q* = 0.3.

The modified Orr model can resolve the problem stated at the top of the previous section. If *p* declines linearly as shown in Figure S2 and *q* = 0.5, the expected number of simple DMIs after *k* = 40 substitutions is 1.2, which is higher than the observed value. Thus, the modified Orr model predicts that most simple DMIs become complex within a single substitution.

### DMI complexification is pervasive in the RNA model

Is there evidence for rampant DMI complexification in the RNA model as predicted under the modified Orr model? Figure 2 shows that inviable double and triple introgressions, which must be based on complex DMIs of order *n* ⩾ 3 and *n* ⩾ 4, respectively, are quite common, as might be expected if more complex inviable introgressions are generated continuously from simpler ones. However, this is not conclusive evidence of complexification (Figure 6B): inviable introgressions of different complexity might simply arise de novo, with more complex ones appearing at a higher rate, as proposed by Orr (1995) (Figure 6A).

**Figure 6.**
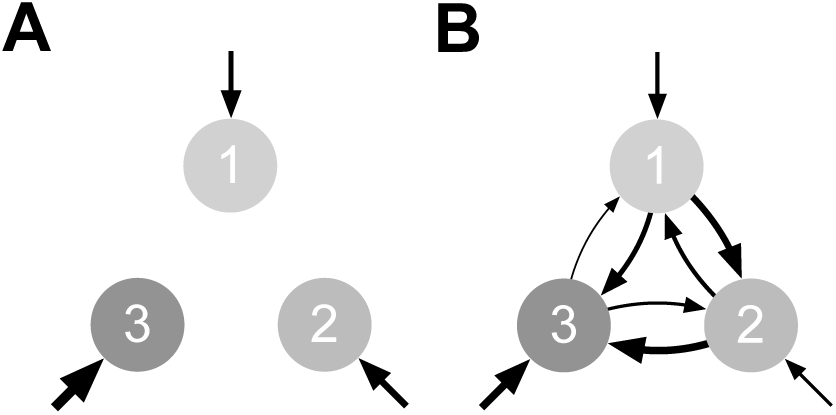
Origins of DMIs of different complexity according to Orr (1995) and this paper. Higher order, *n* (i.e., number of loci involved in a DMI) corresponds to higher complexity. The numbers show *i*, the number of alleles introgressed. An inviable introgression of *i* alleles indicates one or more DMIs of order *n* ⩾ *i* + 1. (A) Orr proposed that DMIs of different complexity arise *de novo*. He also speculated that more complex DMIs arise at higher rates, as indicated by the thickness of the arrows. (B) We propose a “spiraling complexity” model where some DMIs arise *de novo*, but others arise through simplification and complexification of previously existing DMIs. We also propose that complexification is the dominant force, as indicated by the thickness of the 1 → 2 → 3 arrows. The total thickness of the incoming arrows into each value of *i* in (B) is twice that in (A).

To distinguish between the two scenarios, we tracked every inviable introgression in each simulation and classified it according to whether it originated *de novo* or through the simplification or complexification of another inviable introgression present after the previous substitution. Figure 7 shows that initially, most inviable introgressions arise *de novo*. However, after *k* = 40 substitutions, the majority of inviable double and triple introgressions arise through complexification.

**Figure 7.**
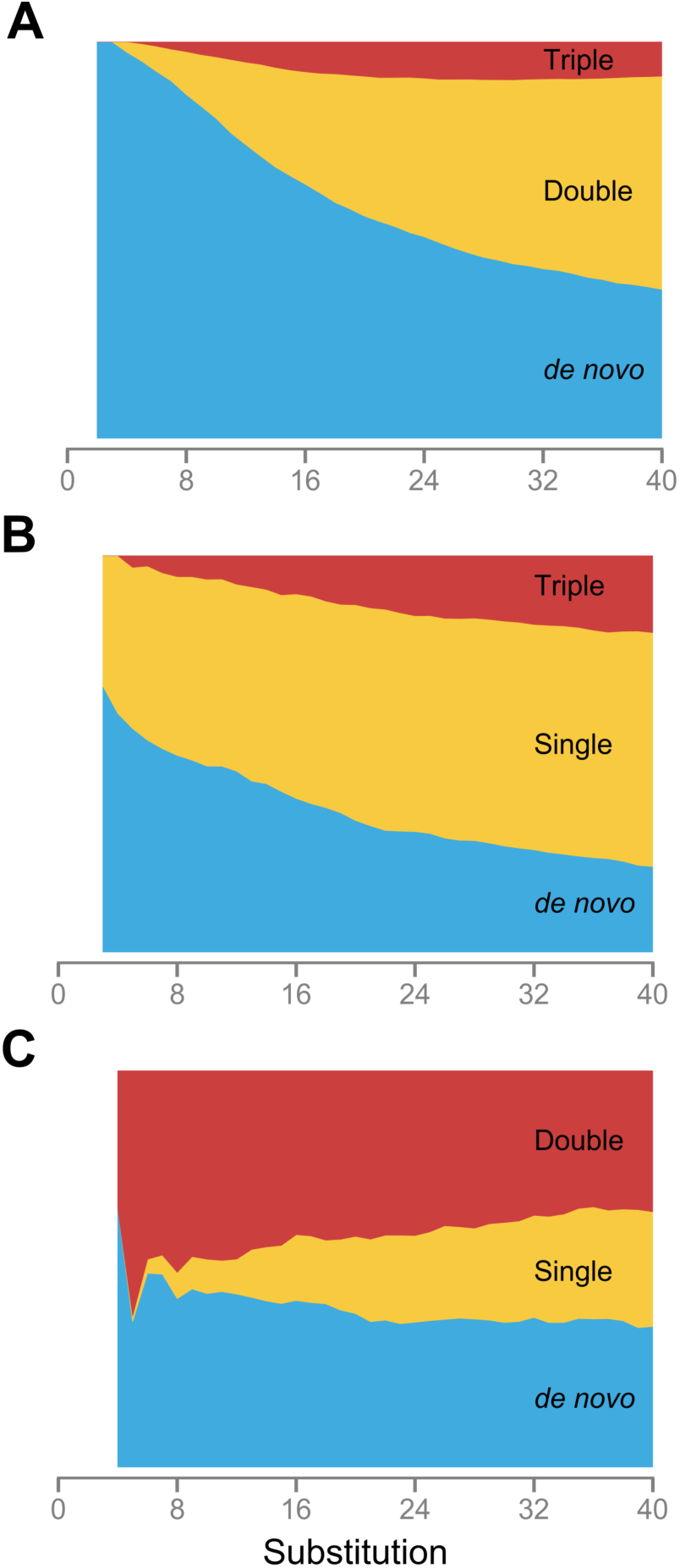
DMI complexification is pervasive in the RNA model. Origin of inviable single (A), double (B), and triple (C) introgressions. The width of each sector indicates the proportion of inviable introgressions arising either *de novo* (blue) or through the modification of another inviable introgression present after the previous substitution. (A) Origin of single introgressions. Incoming arrows into *i* = 1 in Figure 6B: double, 2 → 1; triple,3 → 1. (B) Origin of double introgressions. Incoming arrows into *i* = 2 in Figure 6B: single, 1 → 2; triple, 3 → 2. (C) Origin of triple introgressions. Incoming arrows into *i* = 3 in Figure 6B: single, 1 → 3; double, 2 → 3.

When we compared the final numbers of inviable single and double introgressions we found that they were strongly negatively correlated (Spearman’s rank correlation coefficient: *ρ* = -0.469, *P* < 10^−6^; Figure S3). This result is difficult to explain under the Orr model, because it envisaged that the rates of origination of DMIs of different complexity are independent. However, a negative correlation would be expected if complexification is important and the rates of complexification vary across simulation runs.

### Reproductive isolation does not snowball in the RNA model

Since inviable introgressions snowball in the RNA model, RI would be expected to snowball as well (prediction #3). However, we found that RI showed a kind of inverse snowball effect— a “slowdown” with divergence (Figure 8A). This pattern has been found in many organisms (e.g., Gourbière and Mallet 2010; Giraud and Gourbière 2012). The slowdown was caused by the fact that RI increased slower than linearly with the number of DMIs (Figure 8B). Thus, DMIs did not act independently of each other on RI. One reason for this non-independence is that the total number of inviable introgressions among highly diverged sequences is high enough (Figure 2) that a substantial fraction of individual sites must participate in multiple DMIs (Figure 3).

**Figure 8.**
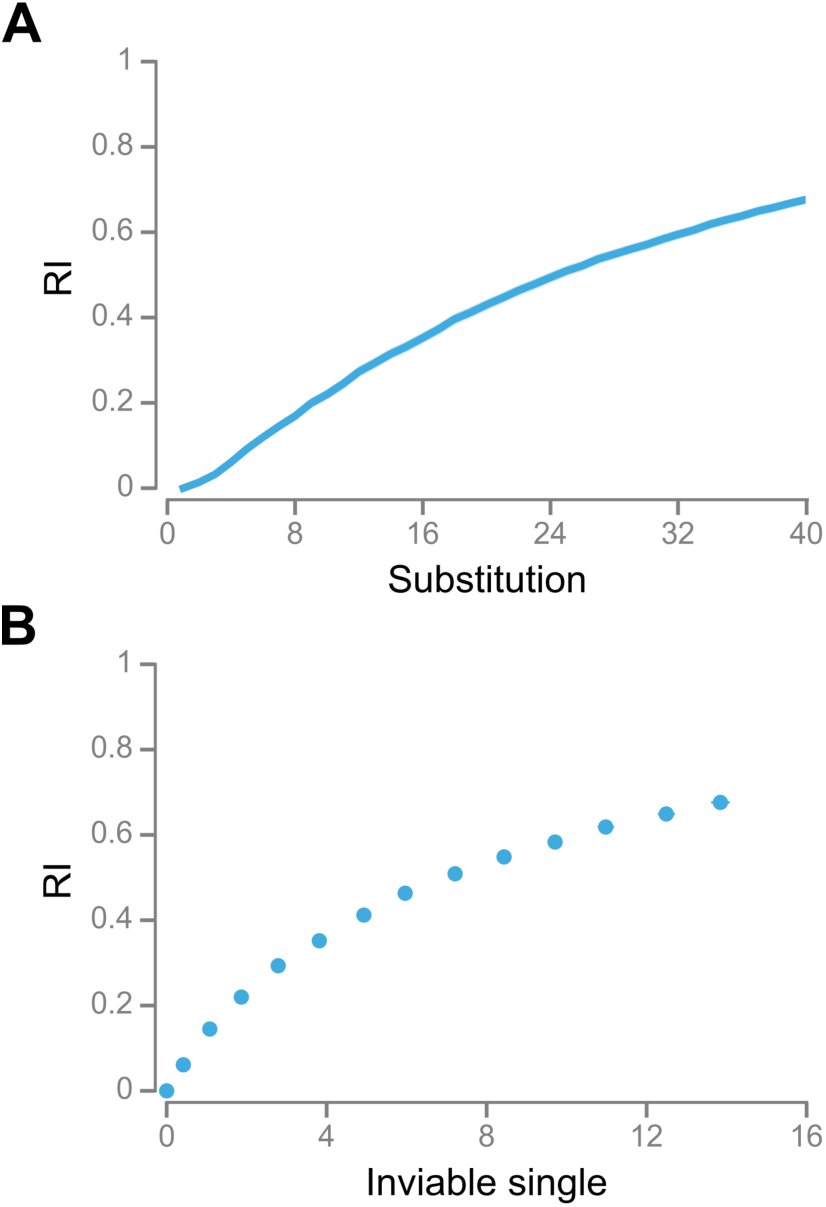
Reproductive isolation (RI) does not snowball in the RNA model. (A) Evolution of RI. Values are means of 10^3^ simulations with *α* = 12. Shaded regions indicate 95% CIs. Mean numbers of inviable single introgressions and RI at *k* = 1, 4, 7,…, 40. Error bars are 95% CIs.

### The structure of the fitness landscape influences the accumulation of DMIs

Figure 9 shows that the accumulation of inviable single introgressions varied extensively between simulations (see Figure S3 for inviable double introgressions). What caused this variation? All simulations took place on the same sequence space, but with different fitness landscapes. Since all fitness landscapes were “holey” (Gavrilets 2004), it follows that the exact pattern of “holeyness” might have had an effect on the evolutionary dynamics. One component of the holeyness of a fitness landscape is the proportion of inviable single mutant neighbors of all the sequences generated during the course of evolution. This measure of the local holeyness of the fitness landscape was strongly positively correlated with the final number of inviable single introgressions (*ρ* = 0.308, *P* < 10^−6^; Figure 9).

**Figure 9.**
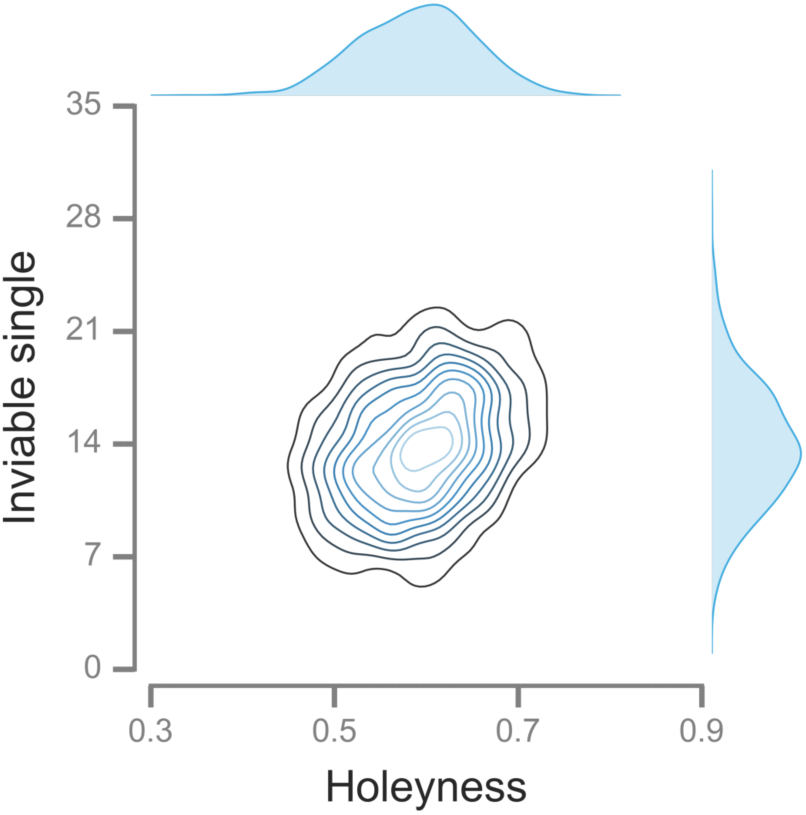
The structure of the fitness landscape influences the accumulation of DMIs in the RNA model. The number of invisable single introgressions accumulated after 40 substitutions is positively correlated with the holeyness of the fitness landscape. One− and two-dimensional kernel density estimates based on 10^3^ simulations with *α* = 12.

What determines holeyness? The fitness landscapes in our RNA folding model have two determinants: the reference structure and the value of *α* (Equation 4). RNA secondary structures can differ in many ways, such as the number and size of base pair stacks, interior loops, and hairpin loops (Schuster *et al.* 1994). The relationship between these structural features and holeyness is difficult to predict *a priori*. For a given reference structure, lower values of *α* are expected to specify fitness landscapes with more inviable sequences (i.e., holes) in them. To evaluate the extent to which these determinants of the fitness landscape influence holeyness, we ran 10^3^ independent evolutionary simulations at each of another four values of α. We found that holeyness was influenced by both determinants of the fitness landscape: it was negatively correlated with *α* (*ρ* = -0.476; *P* < 10^−6^; Figure S4A), and positively correlated with the number of base pairs in the reference sequence, *β*(*ρ* = 0.134; *P* < 10^−6^; Figure S4B). Interestingly, changing the value of *α* had only small effects on the patterns of accumulation of inviable introgressions (Figure S5; Table S3).

## Discussion

We have tested both predictions and assumptions of the Orr model using a computational model of RNA folding. Our results provide mixed support for the snowball effect (Table 2). Inviable introgressions snowballed, but did so more slowly than expected (predictions #1 and #2; Figure 2; Table 1). To elucidate why we tested two assumptions of the Orr model. First, that simple DMIs arise with constant probability, *p* (assumption #1). Although we did detect a decline in *p*, it was not sufficient to account for the pattern of accumulation of inviable single introgressions. Second, we tested assumption #4 that DMIs, once they have arisen, persist indefinitely. We found that this assumption was violated in the RNA model. Instead, DMIs had a tendency to become more complex as further substitutions took place.

**Table 2.** The RNA model provides mixed support for the Orr model.

| Test | Confirmed? | Data |
| --- | --- | --- |
| *Assumption* |  |  |
| 1. $p$ constant with divergence | No. Weak decline. | Figure S2 |
| 2. DMIs have small effects on RI | Yes. | Figure 8A |
| 3. DMIs have independent effects on RI | No. | Figure 8B |
| 4. Simple DMIs persist indefinitely | No. | Figures 4 and 7 |
| *Prediction* |  |  |
| 1. Simple DMIs snowball | No. | Figure 2, Table 1 |
| 2. Complex DMIs snowball | Yes, but slower than expected. | Figure 2, Table 1 |
| 3. RI snowballs | No. | Figure 8A |

Despite the snowballing of inviable introgressions RI did not snowball (prediction #3 of the Orr model) because DMIs did not act independently of each other on RI (assumption #3). These results indicate that RI is a poor indicator for the number of inviable introgressions or DMIs in our model. Thus, the pattern of change in RI with divergence is unsuitable to test the Orr model (Mendelson *et al.* 2004; Johnson 2006; Gourbière and Mallet 2010; Presgraves 2010a). In conclusion, the RNA model provided qualitative support for the central prediction of the Orr model that the total number of DMIs snowballs. However, our results failed to confirm certain predictions of the Orr model, as well as some of its assumptions.

An earlier test of the Orr model using a computational model of gene networks also found no evidence for a snowball effect in RI (prediction #3), and concluded that some assumptions of the Orr model were not met (Palmer and Feldman 2009). However, the extent to which the complexification of DMIs influenced their results is unclear because they did not attempt to investigate the dynamics of the DMIs underlying RI.

In one direct empirical test of the snowball effect, inviable single introgressions affecting pollen sterility were found to accumulate linearly, whereas those affecting seed sterility were found to accumulate faster than linearly (Moyle and Nakazato 2010). Our results suggest a possible explanation for the discrepancy: faster complexification (i.e., higher *q*) of pollen sterility DMIs. Sherman *et al.* (2014) found evidence of greater complexity of the DMIs involved in pollen sterility.

If all DMIs are simple and individual loci are at most involved in one DMI, then the proportion of the fixed differences between species where an allele from one species is deleterious in another species, 𝒫_1_, is expected to increase linearly with genetic distance (Equation 3; Welch 2004). This prediction is contradicted by the observation that 𝒫_1_ is approximately constant over large genetic distances (Kondrashov *et al.* 2002; Kulathinal *et al.* 2004)—a result we call Welch’s paradox. Our results contradict both assumptions behind the prediction that 𝒫_1_ should increase linearly with genetic distance (Welch 2004): most DMIs are complex, and individual loci are involved in multiple DMIs. These effects are expected to act in opposite directions: the former would cause 𝒫_1_ to snowball faster, whereas the latter would cause 𝒫_1_ to snowball more slowly. In the RNA simulations, 𝒫_1_ increased with divergence but did so slower than linearly (Figure S6), indicating that the lack of independence between DMIs dominates the evolution of 𝒫_1_. These results suggest a possible resolution for Welch’s paradox: 𝒫_1_ can be constant even if DMIs snowball if individual loci participate in multiple DMIs. Alternative resolutions of Welch’s paradox have been proposed (e.g., Fraïsse *et al.* 2016).

We found that the vast majority of DMIs in the RNA model were complex. Complex DMIs have been discovered in many introgression studies (reviewed in Wu and Palopoli 1994; Fraïsse *et al.* 2014). For example, Orr and Irving (2001) investigated the sterility of male F1 hybrids between the USA and Bogota subspecies of *D. pseudoobscura* and found that it is caused by an DMI between loci in both chromosomes 2 and 3 of USA and loci in at least three different regions of the X chromosome of Bogota— a DMI of order *n* ⩾ 5. More generally, high-order epistasis appears to be common (Weinreich *et al.* 2013; Kondrashov and Kondrashov 2015; Taylor and Ehrenreich 2015). However, the relative prevalence of simple and complex DMIs in nature is unclear because complex DMIs are more difficult to detect.

Two explanations for the abundance of complex DMIs have been proposed. First, that more complex DMIs evolve more easily than simpler DMIs because they allow a greater proportion of the possible evolutionary paths between the common ancestor and the evolved genotypes containing the DMI (Cabot *et al.* 1994; Orr 1995). Fraïsse *et al.* (2014) tested this mechanism using simulations and concluded that it is unlikely to be effective. Second, that the number of combinations of *n* loci increases with *n* (Orr 1995). This explanation is difficult to evaluate in the absence of more information on the probability of origination of complex DMIs. Our results indicate that that probability could be higher than previously thought because complex DMIs are continuously generated from simpler DMIs. Indeed, our results suggest a new explanation for the abundance of complex DMIs: that DMIs have a tendency to become increasingly complex with divergence.

Our study has identified one determinant of the origination and complexification of DMIs: the holeyness of the fitness landscape. In a holey fitness landscape, our measure of holeyness is inversely related to the mutational robustness of the genotypes assayed (van Nimwegen *et al.* 1999; Ancel and Fontana 2000). In our model (as in Orr’s) “populations” are assumed to contain a single genotype; periodically, a mutant genotype arises and either goes to fixation or disappears. In such a model, mutational robustness is not expected to evolve (van Nimwegen *et al.* 1999). Individual-based simulations would allow us to investigate the intriguing possibility that factors that influence the evolution of mutational robustness (e.g., mutation rate, recombination rate, environmental fluctuations: Ancel and Fontana 2000; Wilke *et al.* 2001; Gardner and Kalinka 2006; Azevedo *et al.* 2006) may influence the accumulation of DMIs.

Perhaps the central insight from our study is that DMIs have a tendency to become more complex. At first glance this claim might seem baffling. Can a DMI really be simple one moment and complex the next? The solution to this puzzle rests, we believe, on the difference between a DMI having a certain order *n* and our ability to *infer* that it has order *n* through genetic crosses. Imagine that one lineage has accumulated 3 consecutive substitutions at loci A, B, and C. Derived alleles are indicated by uppercase letters, and ancestral alleles by lowercase letters. Now, imagine that there is a complex DMI of order *n* = 3 between the alleles *a*, B, and *c*, and that there are no simple DMIs between any pairs of the three alleles (i.e., *a*/*B*, *a*/*c*, and *B*/*c*). For simplicity, we also assume that none of the other alleles at the A, B and C loci are involved in DMIs. The existence of a DMI is defined in the strict sense that any conceivable genotype containing all alleles involved in the DMI is inviable (conversely, the absence of a DMI indicates that at least one of the genotypes containing all alleles involved in the DMI are viable). Despite the *a*/*B*/*c* DMI being complex, after two substitutions, our introgression and rescue tests (File S2) would detect a nonexistent simple DMI between alleles *a*, and *B*. The actual complex DMI can only be inferred after the third substitution has taken place. In the language we have been using so far, the simple DMI would appear to become more complex.

We believe that our central finding that DMIs have a tendency to become more complex is independent of the details of our model. Other results, such as the precise rate of accumulation of DMIs or inviable introgressions, are likely to be influenced by the details of our model. The extent to which the RNA folding model is representative of other types of epistatic interactions (e.g., in gene networks) is unclear. One possible criticism is that we used very short sequences and that these are likely to experience unusually strong epistatic interactions. Orr and Turelli (2001) estimated *p* ≈ 10^−7^ in *Drosophila*, and Moyle and Nakazato (2010) estimated *p* ≈ 10^−9^ in *Solanum*, much lower values than found in our simulations. However, an evolution experiment in *S. cerevisiae* detected a simple DMI between two lineages that had only accumulated 6 unique mutations each (*k* = 12) (Anderson *et al.* 2010). This indicates a value of *p* ≈ 0.015, remarkably close to what we estimated in the RNA model (Figure S2). Our approach to testing the Orr model can be applied to other computational models of biological systems, such as, transcription-factor binding (Tulchinsky *et al.* 2014; Khatri and Goldstein 2015), gene networks (ten Tusscher and Hogeweg 2009; Palmer and Feldman 2009), and metabolic networks (Barve and Wagner 2013).

Our results were robust to a broad range of holey fitness landscapes defined in the RNA folding model. However, the holey landscape model makes two strong assumptions about the fitness landscape: all viable genotypes had the same fitness, and all low fitness genotypes were completely inviable. Neither assumption is met universally: many alleles involved in DMIs appear to have experienced positive selection during their evolutionary history (Presgraves 2010b; Rieseberg and Blackman 2010; Maheshwari and Barbash 2011), and some DMIs are only mildly deleterious rather than lethal (Presgraves 2003; Schumer *et al.* 2014). These assumptions can be relaxed in the RNA folding model (e.g., Cowperthwaite *et al.* 2005; Draghi *et al.* 2011) and in other models (e.g., Palmer and Feldman 2009; Tulchinsky *et al.* 2014; Khatri and Goldstein 2015).

Studies like ours can test whether the snowball effect occurs under well-defined circumstances. However, they cannot test the *reality* of the snowball effect; introgression studies with real organisms remain the only way to do so (Matute *et al.* 2010; Moyle and Nakazato 2010; Matute and Gavin-Smyth 2014; Sherman *et al.* 2014; Wang *et al.* 2015).

## Acknowledgments

Tim Cooper, Tiago Paixão, Leonie Moyle, and two anonymous reviewers provided useful comments on the manuscript. We had helpful discussions with Rafael Guerrero, Peter Olofsson, and Jeff Tabor. We used the Maxwell and Opuntia clusters from the Center of Advanced Computing and Data Systems (CACDS) at the University of Houston. CACDS staff provided technical support. The National Science Foundation (grant DEB-1354952 awarded to R.B.R.A.) funded this work.

